# Loss of mutant TP53 does not impair the sustained proliferation, survival or metastasis of diverse cancer cells

**DOI:** 10.1101/2020.03.30.015263

**Authors:** Zilu Wang, François Vaillant, Catherine Chang, Chris Riffkin, Elizabeth Lieschke, Sarah Diepstraten, David CS Huang, Jane E Visvader, Marco J Herold, Gemma L Kelly, Andreas Strasser

## Abstract

The tumour suppressor *TP53* is the most frequently mutated gene in human cancer and these aberrations confer poor chemotherapeutic responses^1-3^. Point mutations typically cluster in the DNA binding domain, with certain ‘hot-spot’ residues disproportionally represented^1-4^ These mutations abrogate binding of the TP53 transcription factor to DNA and thereby prevent upregulation of genes critical for tumour suppression (loss-of-function)^1-3^. Mutant TP53 is reported to additionally contribute to tumour development, sustained growth and metastasis not only through dominant-negative effects on wild-type TP53^5^ but also through neomorphic gain-of-function (GOF) activities^6^. Understanding the contributions of these postulated attributes of mutant TP53 to the development and expansion of tumours will facilitate the design of rational therapeutic strategies. Here we used CRISPR/CAS9 to delete mutant *TP53* in a panel of diverse human cancer cell lines. The loss of mutant *TP53* expression had no impact on the survival, proliferative capacity or metabolic state of the tumour cells, nor did it sensitise them to cellular stresses and chemotherapeutic agents. These data suggest that putative GOF effects of mutant TP53 are not universally required for the sustained survival and proliferation of fully malignant cells. Therefore, therapeutic approaches that abrogate expression or function of mutant TP53 would not be expected to have substantial impact.

---

TP53 is a transcription factor that acts as a potent tumour suppressor through oncogenic stress induced upregulation of genes involved in diverse cellular responses, including apoptotic cell death, cell cycle arrest, cellular senescence and DNA repair^7,8^. Accordingly, aberrations in the gene encoding TP53 are found in diverse human cancers with an overall frequency of ∼50%^2^. While some of these aberrations cause complete loss of the TP53 protein (modelled experimentally by *TP53* knockout mice), most mutations selected for in human cancers result in substitution of single amino acids, usually in the DNA binding domain, and high level expression of stabilised mutant TP53 proteins is a common feature of malignant cells carrying *TP53* mutations^1,9^. Mutant TP53 proteins have been proposed to drive malignant transformation and to sustain tumour growth via three processes that are not mutually exclusive: (1) loss-of-function (LOF), i.e. an inability of mutant TP53 to activate expression of the genes that are transcriptionally activated by wild-type (wt) TP53 protein to induce cell cycle arrest, apoptotic cell death or other cellular responses; (2) dominant-negative effects (DNE), i.e. mutant TP53 represses wt TP53 function through engagement in mixed mutant TP53/wt TP53 tetramers; and (3) gain-of-function (GOF) effects (Fig 1a) ^1-3,6^. The GOF effects are thought to be mediated through neomorphic interactions of mutant TP53 protein with signal transducers or transcription factors activating cellular responses that are not controlled by wt TP53^4^. The LOF and DNE of mutant TP53 are widely accepted to be critical for malignant transformation, with the importance of the DNE restricted to early stages of tumorigenesis when nascent neoplastic cells still express wt TP53 that is functionally repressed by mutant TP53^5,10,11^. Sustained LOF was proven to be critical for continued tumour growth using mouse models in which expression of wt TP53 could first be switched off to drive tumorigenesis but then restored again in the malignant cells, resulting in suppression of tumour expansion through induction of either apoptotic death or cell cycle arrest/senescence, depending on the tumour cell type^12-14^. It is less clear how important the GOF effects of mutant TP53 are for tumour initiation and sustained tumour growth. Resolving this issue has important ramifications for the design of novel cancer therapies. Genetic removal or pharmacological inhibition of mutant KRAS diminishes the growth of cancers that are driven by such mutations^15^. If the GOF effects of mutant TP53 were likewise essential for sustained tumour growth, drugs that could abolish the expression or function of mutant TP53, for example ones based on PROTAC technology^16,17^, would be predicted to have substantial therapeutic benefit. Such a therapeutic benefit resulting directly from loss of the GOF activity of mutant TP53 was reported in a study using knock-in mice expressing a hot-spot mutant TP53, R248Q, where its expression could be extinguished in the resulting lymphomas through CRE-mediated mutant *TP53* gene deletion^18^. The loss of mutant TP53 expression curbed the growth of the tumours, albeit to a modest extent. These experiments supported a critical role for the GOF activity of mutant TP53 in sustaining tumour expansion since this manipulation would generate TP53-deficient malignant cells.

**Fig. 1:**
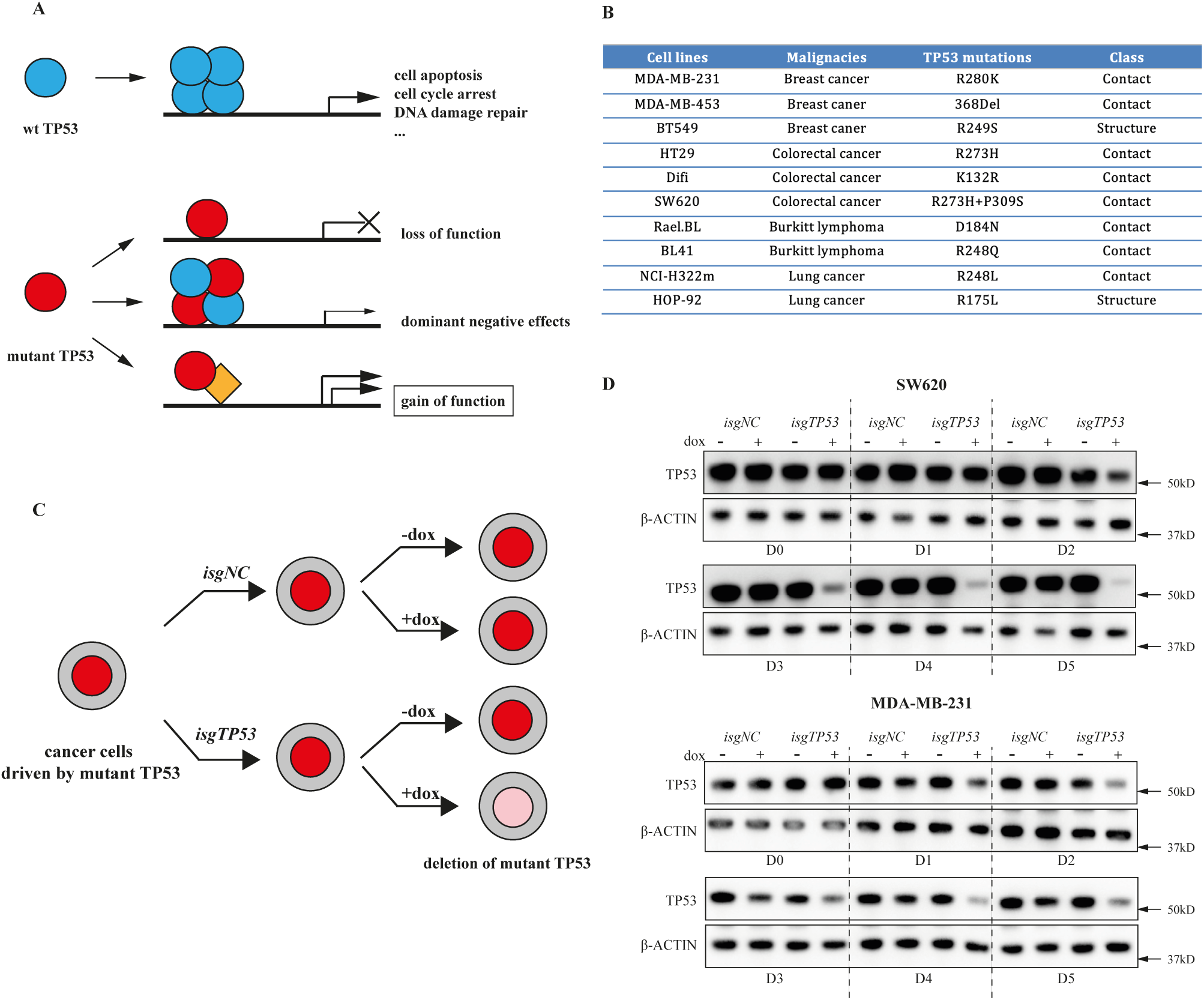
Progressive generation of TP53-null variants of human cancer cell lines that are driven by mutant TP53 using CRISPR/CAS9 technology. **a**. Schematic of how wt TP53 functions as a tumour suppressor and mechanisms by which mutant TP53 is postulated to contribute to tumorigenesis. **b**. Table summarizing the name and cellular origin of the selected mutant TP53 harbouring human cell lines examined, with the TP53 mutations detailed. **c**. Schematic to illustrate the proposed strategy for progressively removing mutant TP53 expression using CRISPR/CAS9 methodology in the selected human cancer-derived cell lines. **d**. Western blotting to demonstrate the progressive loss of mutant TP53 protein in the human cancer cell lines SW620 and MDA-MB-231 infected with the sgRNAs targeting TP53 over time. The same cells transduced with a NT sgRNA are shown for comparison. Probing for β-ACTIN was used as a protein loading control.

## Inducible loss of mutant TP53 does not reduce proliferation or survival of diverse human cancer cell lines

The topical large investments by pharmaceutical and biotechnology companies into the development of drugs that abrogate mutant TP53 expression or function, ignited us to examine the requirement for mutant TP53 expression for the sustained growth of an extensive panel of human cancer-derived cell lines using an inducible CRISPR/CAS9 platform^19^. Mutant TP53 was deleted in an inducible manner in ten cell lines containing 11 different TP53 mutations derived from four types of human cancers, namely breast, colorectal and lung carcinomas as well as lymphomas (Fig. 1b). These tumour cells were transduced with a vector for stable expression of Cas9 plus doxycycline (dox) inducible vectors for the expression of a mutant *TP53*-specific guide RNA (*isgTP53*) or a control guide RNA (*isgNC*). The transduced tumour cells were then treated with dox (to induce sgRNA expression) or left untreated and their survival and proliferation were monitored from day 0 to day 5 of treatment (Fig. 1c). The progressive removal of mutant TP53 protein was verified by Western blotting (Fig. 1d and Extended Data Fig. 1). Inducible loss of mutant TP53 had no impact on the *in vitro* survival or proliferation of all cancer cell lines tested (Fig. 2a,b, Extended Data Fig. 2a) Flow cytometric analysis confirmed that deletion of mutant TP53 had no impact on cell cycling (Fig. 2c, Extended Data Fig. 2b). Since these experiments were conducted during and immediately after the removal of mutant TP53, it appears unlikely that acquisition of mutations that provide compensation for the loss of mutant TP53 would explain these findings.

**Fig. 2:**
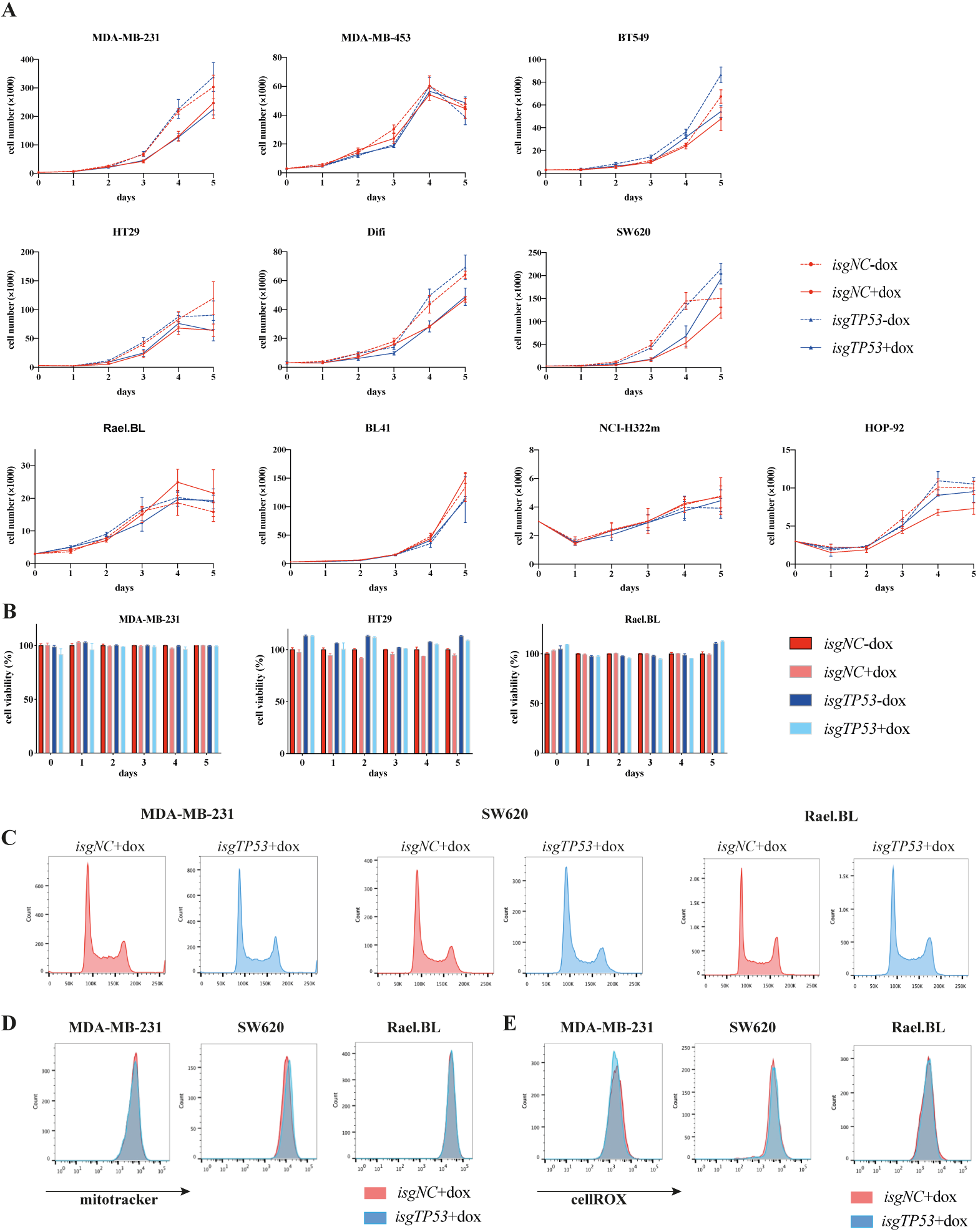
Loss of mutant TP53 does not impair the sustained cell proliferation, survival, mitochondrial content and ROS levels in human cancer cell lines. **a**. *In vitro* growth of the indicated human cancer cell lines with or without dox mediated induction of a mutant *TP53* specific or a control sgRNA. **b**. *In vitro* survival of the cells described in (**a**). **c**. Cell cycle analysis of the cells described in (**a**). **d**. Mitotracker staining of the cells described in (**a**). **e**. CellRox staining of the cells described in (**a**). The analyses described in (**c**), (**d**) and (**e**) were conducted 2 days after the cells had been treated with dox for 5 days (see (**a**)). Data in (**a**) and (**b**) are presented as mean±SD of experiments conducted in triplicates.

### Inducible loss of mutant TP53 does not impact mitochondrial activity or ROS levels in human cancer cell lines

Adaptation of cell metabolism (i.e. the Warburg effect^20^) is considered one of the hallmarks of cancer^21^ and high intracellular levels of reactive oxygen species (ROS) levels are often associated with increased cellular metabolism^22^. Mutant TP53 has been reported to regulate metabolic changes and impact the levels of ROS in cancer cells^23^. Mitotracker and CellROX staining revealed that loss of mutant TP53 had no impact on mitochondrial content and activity, or on the intracellular levels of ROS, respectively, in any of the cancer cell lines tested (Fig. 2d, e, Extended Data Fig. 2c, d). These data show that continuous expression of mutant TP53 is not required for the sustained metabolic adaptation of cancer cells.

### Loss of mutant TP53 does not impact response to nutrient deprivation or cytotoxic drugs in human cancer cell lines

The ability of cancer cells to tolerate diverse stresses, such as deprivation of nutrients and growth factors, is critical for ensuring the survival and continual expansion of the tumour^24,25^. Of note, the GOF effects of mutant TP53 have been reported to assist malignant cells in dealing with and adapting to stress^26,27^. To determine the importance of the GOF effects of mutant TP53 on cancer cells under stress, we cultured the parental and TP53-depleted cancer cell lines in medium containing only 3% or 1% foetal calf serum (FCS) (Figure 3a-e and Extended Data Fig. 3a-d) since nutrient deprivation can induce cellular stress and metabolic reprogramming in malignant cells^28^. Culturing in 1% FCS plus dox induction could induce significant death in both the parental and mutant TP53-depleted SW620 (∼50%) and HOP-92 cells (∼20%), but this was not seen in the other cell lines (Fig. 3b, Extended Data Fig. 3a). This is likely due to the sensitivity of these cells to dox toxicity when cultured in low serum. By contrast, when cells were cultured in 3% FCS, none of the cell lines displayed significant cell death following the deletion of mutant TP53 (Extended Data Fig. 4b). Furthermore, we could not detect any impact of induced loss of mutant TP53 on cell growth, cell cycling, mitochondrial content and activity or ROS levels in any of the cell lines tested, both in medium containing 1% or 3% FCS (Fig. 3a,c-e, Extended Data Fig. 3b-d, Fig. 4a and Fig. 5a-c). Mutant TP53 has been reported to render malignant cells resistant to diverse anti-cancer drugs^29^. However, we found that deletion of mutant TP53 did not significantly increase chemotherapeutic drug induced killing of the cancer cell lines tested (Extended Data Fig. 6). These results demonstrate that mutant TP53 does not have a marked impact on the ability of cancer cells to adapt to conditions of stress, such as nutrient deprivation or treatment with anti-cancer agents.

**Fig. 3:**
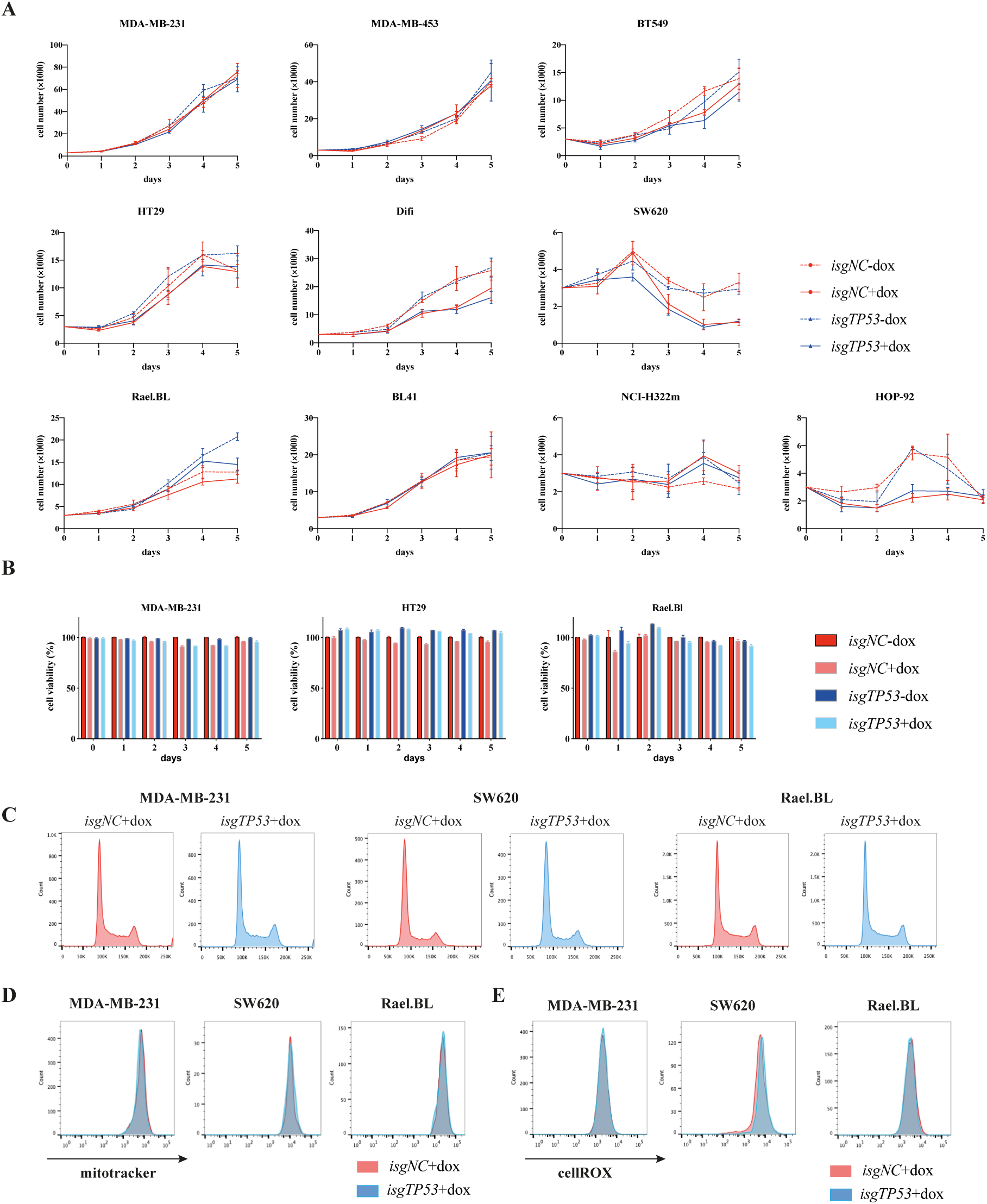
Loss of mutant TP53 does not impair the ability of cancer cells to adapt to conditions of stress. **a**. *In vitro* growth of the indicated human cancer cell lines with or without dox mediated induction of a mutant *TP53* specific or a control sgRNA grown in medium with 1% FCS. **b**. *In vitro* survival of the cells described in (**a**). **c**. Cell cycle analysis of the cells described in (**a**). **d**. Mitotracker staining of the cells described in a. **e**. CellRox staining of the cells described in (**a**). The analyses described in (**c**), (**d**) and (**e**) were conducted after 2 days in culture in medium with 1% FCS, which was done after the cells had been treated with dox for 5 days in normal medium with 10% FCS. Data in (**a**) and (**b**) are presented as mean±SD of experiments conducted in triplicates.

**Fig. 4:**
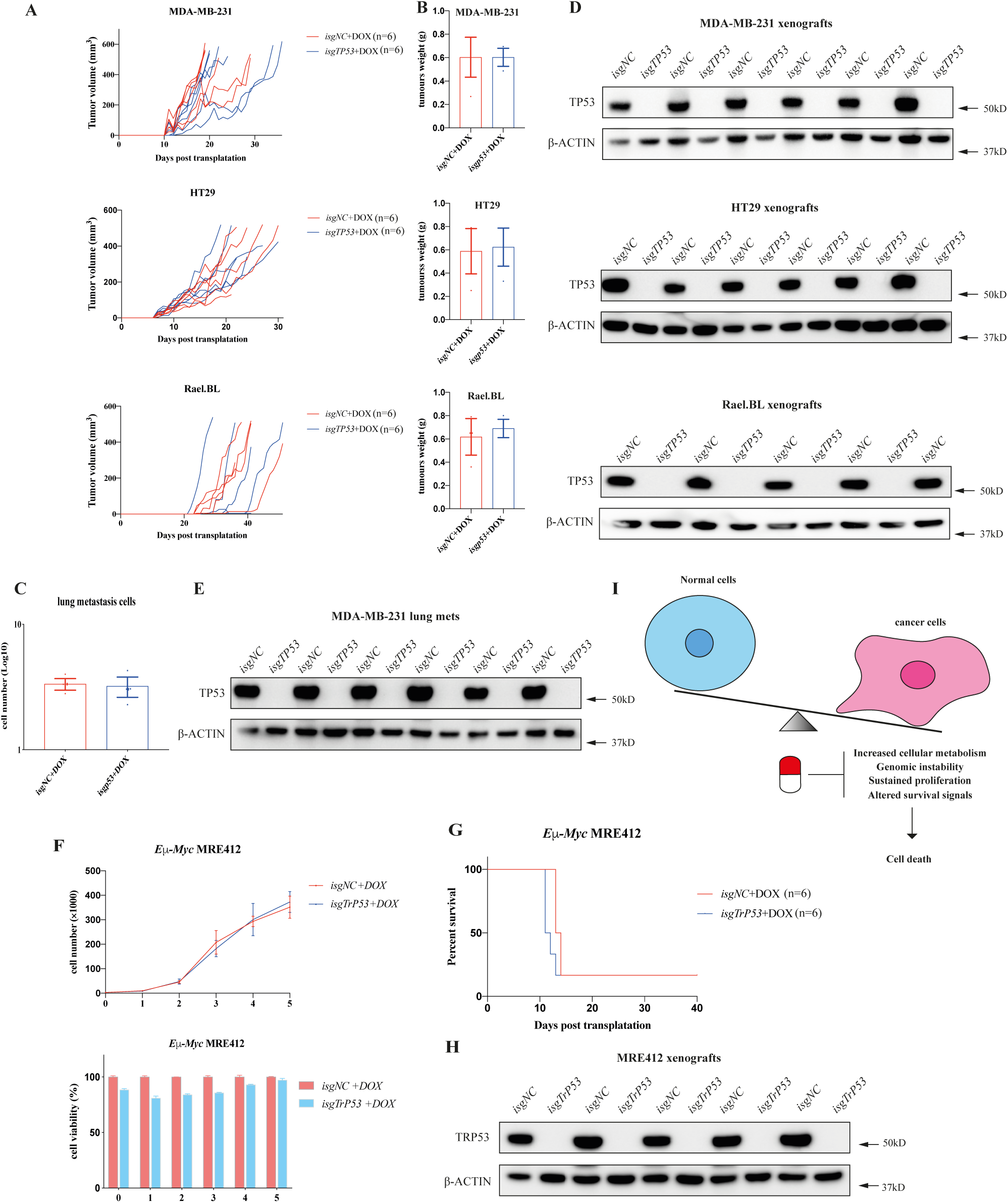
Loss of mutant TP53 does not impair tumour growth and metastasis *in vivo*. **a**. Growth of the human cancer cell lines MDA-MB-231, SW620 and Rael-BL, either parental or their mutant TP53-deficient derivatives, in NSG mice (n=6 mice per cell line). **b**. Weights of the tumours from (**a**) at the ethical endpoint. **c**. Numbers of metastatic cells in the lungs of NSG mice that had been injected with MDA-MB- 231 breast cancer cells, either parental or their mutant TP53-deficient derivatives, into their mammary fat pads (n=6 mice per cell line). **d**. Western blot analysis of the tumours from (**a**) to verify the presence of mutant TP53 protein in the parental cells and to confirm the absence of mutant TP53 protein in the tumours arising from the injected cells that had been engineered with sgRNAs targeting TP53. Probing for β-ACTIN was used as a protein loading control. **e**. Western blot analysis of the metastases from (**c**) to verify the presence of mutant TP53. Probing for β-ACTIN was used as a protein loading control. **f**. *In vitro* survival and growth of the *Eμ-Myc* lymphoma cell line, MRE412, either the parental line or its mutant TP53 depleted derivative (n=6 mice per cell line). **g**. Kaplan-Meier survival curves of mice transplanted with the parental *Eμ-Myc* lymphoma cell line, MRE412, or its mutant TP53 depleted derivative line from (**f**). **h**. Western blot analysis of the tumours from (**g**) to verify the presence of mutant TP53 in the parental cells or its absence in their mutant TP53-deficient derivatives. Probing for β-ACTIN was used as a protein loading control. **i**. Possible strategies for the treatment of cancers with mutant TP53 or loss of TP53. Data in (**b**), (**c**) and (**f**) are presented as mean±SD of experiments conducted in triplicates.

### Clonal populations of mutant TP53-deficient human cancer cell lines behave similarly to their parental mutant TP53 expressing counterparts

A potential caveat of the data presented thus far is that within the polyclonal populations of cancer cell lines that had been induced to express the mutant TP53 targeting sgRNA, a small amount of mutant TP53 protein remained detectable by Western blot analysis (Fig. 1d, Extended Data Fig. 1). To determine whether the residual, albeit markedly reduced levels of mutant TP53, could have impacted the read-out of the experiments, single cell derived clones (2-3 per line) lacking detectable mutant TP53 protein were established from representative cancer cell lines: MDA-MB-231 (breast cancer, R280K), HT29 (colorectal cancer, R273H) and Rael-BL (Burkitt lymphoma, D184N) (Extended Data Fig. 7a, 8a, 9a). Importantly, the complete loss of mutant TP53 had no impact on the proliferation, survival, cycling, mitochondrial content and activity, ROS levels and response of these cell lines to anti-cancer agents, even in medium containing only 3% or 1% FCS (Extended Data Fig7b-g, 8b-g, 9b-g).

### Loss of mutant TP53 does not reduce growth or metastasis of human cancer cell lines *in vivo*

Of note, MDA-MB-231, HT29 and Rael-BL cancer cell line derived clones with complete loss of mutant TP53 were able to grow in immune-deficient NSG mice in a manner comparable to their parental counterparts (Fig. 4a), exhibiting similar tumour weights at the ethical endpoint (Fig. 4b). For MDA-MB-231 breast cancer cells implanted into mammary fat pads, metastasis was analysed but no significant differences in the numbers of lung metastases were observed between the mutant TP53 deleted cells *vs* their parental counterparts (Fig. 4c). Western blotting confirmed the loss of mutant TP53 in the primary breast tumours as well as in the lung metastases (Fig. 4d, e). Inducible loss of mutant TRP53 has been shown to induce apoptosis in mouse lymphomas and inhibit their growth *in vivo*^18^. In contrast, we found that complete removal of mutant TRP53 had no impact on the *in vitro* survival or proliferation of the *Eμ-Myc* lymphoma cell line MRE412, that expresses mutant TRP53 (Fig. 4f), or its ability to grow in mice (Fig. 4g). Flow cytometric analysis of GFP and Ly5.2 expression verified that the malignant cells were derived from the transplanted *Eμ-Myc* lymphoma cells (Extended Data Fig. 10), and loss of mutant TRP53 was confirmed by Western blotting (Fig. 4h). Our findings are consistent with the observations that tumours from TP53-deficient (*TP53-/-*) mice, the state induced by genetic loss of mutant TP53 in cancer cells, are highly aggressive^30,31^ and markedly resistant to diverse anti-cancer agents^32,33^. Collectively, these results demonstrate that mutant TP53 is not critical for the sustained *in vivo* growth or metastasis (the latter only tested for MDA-MB-231 cells) of a diverse set of human cancer-derived cell lines, as well as a mouse lymphoma-derived cell line.

### shRNA-mediated reduction of mutant TP53 also does not impair the survival and proliferation of several human cancer cell lines

In contrast to our findings, some previous studies have shown that reduction of mutant TP53 expression using RNA interference was able inhibit the *in vitro* growth and kill certain human cancer cell lines^34,35^. To try to reconcile these studies with our findings, the previously published *TP53* targeting (*shTP53*) and control (*shNC*) shRNA sequences were cloned into a dox-inducible viral? vector and transduced into the same cancer cell lines used in those studies: SKBR3, HUH-7 and T47D. The reduction in mutant TP53 levels (Extended Data Fig. 11a) as well as cell survival and proliferation were monitored from day 0 to day 5 after dox induction of the shRNAs. Dox induction caused minor cell death and delayed cell proliferation not only in the cells transduced with *ishTP53* but also those transduced with the control *ishNC* in all three cell lines tested (Extended Data Fig. 11b). Of note, dox induced expression of both *ishTP53* and *ishNC* also induced some death and inhibited the *in vitro* growth of the HT29 cancer cell line that had previously already been depleted for mutant TP53 using CRISPR, but not in the other two cell lines, MDA-MB-231 and SW620 (Extended Data Fig. 11c). This indicates that these shRNAs can exert substantial non-specific toxicity. Of note, a recent report found that several novel targets (e.g. PIM1, MAPK14) for cancer therapy that had been identified using RNA interference could not be validated using CRISPR technology and that the killing of the cancer cell lines induced by the shRNAs was in fact due to the targeting of other proteins^36^.

## Discussion

Collectively, our studies using 10 cell lines representing four different types of human cancers and one mouse lymphoma line, overall representing twelve different mutants of TP53, reveal that the GOF effects of mutant TP53 are not universally required for the sustained survival and proliferation of malignant cells. This suggests that drugs capable of abrogating expression or function of mutant TP53, for example based on PROTAC technology^16^, would be predicted to not afford substantial therapeutic benefit. Perhaps drugs that are reported to restore wt TP53 activity in at least certain mutant TP53 proteins, such as PRIMA-1 or its methylated analogue APR-246^37^ may have greater impact in such cancers, although it must be noted that these compounds have also been reported to kill malignant cells in a manner independent of the presence of mutant or wt TP53^38,39^. An alternative approach to improve the current poor outlook for patients with mutant TP53 cancers would be to identify dependencies (i.e. vulnerabilities) driven by mutant TP53 (or loss of TP53) that do not manifest in non-transformed cells and hence may be exploited by using already existing or novel therapeutics (Fig. 4i).

## Supporting information

supplementary material combined

## Materials and Methods

### Cells

Burkitt lymphoma cell lines, BL41 and Rael-BL were cultured in RMPI1640 medium (Thermofisher) supplemented with 10% FCS (Sigma), 2mM L-glutamine (Thermofisher), 1mM sodium pyruvate (Thermofisher) and 50 μM α-thioglycerol (Sigma). SW620, MDA-MB-231 and MDA-MB-453 were maintained in L-15 medium (Thermofisher) supplemented with 10% FCS. HOP-92, NCI-H322m, Difi, HT29 and BT549 cell lines were cultured in RPMI1640 medium with 10% FCS. The primary *Eμ-Myc* lymphoma cells were separated from the lymphoid organs of *Eμ-Myc* transgenic mice and cultured in DME medium (Thermofisher) supplemented with 10% FCS, 50μM β-mercaptoethanol (Sigma) and 100μM Asparagine (Sigma). The authenticity of all cell lines was verified by genomic analyses.

### Virus packaging and infection of cell lines

The construction of the inducible sgRNA and shRNA expression vectors and virus production were performed as previously described^19^. A negative control sgRNA (*isgNC*) which targeted mouse *Bim* exon 2 (5’-GACAATTGCAGCCTGCTGAG) and sgRNA against human *TP53* exon 5 (5’- GAGCGCTGCTCAGATAGCGA) were used. For adherent tumour cell lines, 5×10^5^ cells were plated into 6-well plates along with 3 mL viral supernatant and cultured overnight. For suspension tumour cell lines, 1×10^5^ cells were suspended in 5 mL viral supernatant and centrifuged at 2200 rpm for 2 h at 32°C. GFP (indication of sgRNA) and mCherry (marker of CAS9 expression) double positive cells were sorted for subsequent experiments.

### Western blotting

Total protein extracts were obtained from cultured tumour cells and from tumours of mice by using RIPA buffer. A total of 20 μg protein was loaded onto 10% Bis-Tris gels (Thermo Fisher) for separation by electrophoresis and transferred to nitrocellulose membranes. For probing we used antibodies against β-Actin (clone AC-74, Sigma), TP53 (clone DO-7, Santa Cruz) and TrP53 (clone CM5, Leica biosystems) and an anti-mouse/rabbit secondary antibody (Southern Biotech). Protein bands were visualised by Western HRP substrate (Millipore) and ChemiDoc imaging system (Bio-rad).

### Cell counting

3×10^3^ cells were plated into 96-well flat bottom plates in medium containing different concentrations of FCS, in the presence or absence of dox (10 μg/mL) to induce expression of sgRNAs. From day 0 to day 5 after treatment with dox (or no addition of dox), cell numbers were determined: 90 μL cell culture was mixed with 10 μl APC- conjugated beads (concentration known) and 2.5 μg/mL Propidium Iodide (PI), followed by flow cytometric analysis using an LSR IIW (BD). Calculation of cell numbers = 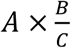 where A = number of PI negative cell events; B = assigned bead counts; C = number of beads events.

### Viability assessment

3×10^3^ cells were plated into 96-well flat bottom plates in medium containing different concentrations of FCS, with the induction of dox treatment of without it. The cell viability was monitored from day 0 to day 5 after dox induction (or no treatment of dox). For etoposide treatment, 1×10^4^ cells were plated into 96-well flat bottom plates in medium containing 10% FCS. 24h after seeding, the cells were incubated with a serial dilution of etoposide for 48h. Cells were stained with 2.5 μg/mL PI and the cell survival was analysed by flow cytometric analysis.

### Cell cycle analysis

1×10^5^ cells were plated into 24-well plates with medium containing 1% FCS, 3% FCS or 10% FCS. After 48 h, cells were fixed using the eBioscience fixation/permeabilisation buffer (Thermo Fisher) and stained with DAPI. DNA content and cell cycle status were analysed by flow cytometry and FlowJo software.

### MitoTracker and CellROX staining

1×10^4^ cells were plated into 96-well flat bottom plates with medium containing 1%, 3% or 10% FCS, respectively. After 48 h, the medium was removed and pre-warmed 200 nM MitoTracker or 5 μM CellROX was added. After incubation for 30 min under growth conditions, cells were harvested for flow cytometric analysis.

### Mice and xenograft transplants

All experiments with mice were carried out in accordance with both the guidelines of the Melbourne Directorate Animal Ethics Committee and the Walter and Eliza Hall Institute of Medical Research Ethics Committee. 6-7 week-old NSG female mice were obtained from The Walter and Eliza Hall Institute’s breeding facility. For the human cancer cells lines MDA-MB-231, SW620 and Rael-BL, the parental cells and their corresponding mutant TP53-depleted derivatives were transplanted (2×10^6^ tumour cells per injection for MDA-MB-231 and HT29 cell lines, 4×10^6^ tumour cells per injection for Rael.BL cell line) subcutaneously into the flank of mice. Tumour size was monitored until they had reached the mandated animal ethics endpoint. To examine metastasis, MDA-MB-231 cells were injected into mammary fat pads. When the primary MDA-MB-231 tumours had grown to ethical endpoint (600 mm^3^), the lungs of tumour burdened mice were excised and single cell suspensions were generated. GFP/mCherry double positive cells were sorted for further analysis.

*Eu-Myc* lymphoma cell lines (C57BL/6J-Ly5.2 background), MRE412, both the parental cells and their corresponding mutant TRP53-depleted derivatives were injected intravenously into C57BL/6J-Ly5.1 males that had been purchased from the Walter and Eliza Hall Institute’s breeding facility.

### Statistical analyses

GraphPad Prism was used for statistical analysis. Error bars indicated the standard deviation of 3 independent experiments. Two-tailed *t* tests were used to compared between 2 data sets.

## Data Availability Statement

Data and reagents are available on request from A.S.

## Acknowledgements

The authors thank Giovanni Siciliano, Dan Fayle, Louise Spencer and Eren Loza for mouse husbandry; S. Monard and his team at the WEHI Flow Cytometry Unit for their assistance. This work was supported by funding from: the Victorian Cancer Agency Fellowship (MCRF 17028) awarded to G.L.K, Cancer Council Victoria, grants-in-aid #1086157 and #1147328 awarded to G.L.K.; the National Health and Medical Research Council, Project Grant #1086291 awarded to GLK, Program Grant #101671 awarded to A.S. and J.E.V., Fellowship #1020363 awarded to A.S., Fellowship #1102742 awarded to J.E.V.; the Leukaemia Foundation Australia grant awarded to G.L.K and A.S., the Leukaemia and Lymphoma Society Grant #7001-13, awarded to A.S.; the estate of Anthony (Toni) Redstone OAM and The Craig Perkins Cancer Research Foundation; and operational infrastructure grants through the Australian Government NHMRCS IRIISS and the Victorian State Government Operational Infrastructure Support.

## Author Contributions

A.S.,G.L.K. and Z.W. conceived the project. D.C.S.H. and C.R. provided human cancer cell lines. C.C. assisted with experiments. F.V. conducted and J.E.V. supervised the *in vivo* metastasis experiments. M.J.H supervised the inducible deletion of mutant TP53 in cancer cells. Z.W. performed the cell survival, cell growth, MitoTracker and ROS staining. E.L. and Z.W. performed the cell cycle analysis. S.D and Z.W established the loss of mutant TP53 cell lines. A.S., G.L.K. and Z.W. wrote the paper, which was edited by all other co-authors.

