## supplementary material combined for "Loss of mutant TP53 does not impair the sustained proliferation, survival or metastasis of diverse cancer cells"

**Zilu Wang et al, Extended Data Figures and Tables with Legends**

### Extended Data Fig. 1 Loss of mutant TP53 in human cancer cell lines

Western blot analyses confirming CRISPR/CAS9-mediated deletion of mutant TP53 in the indicated human cancer cell lines, in which a mutant *TP53* specific sgRNA had been induced by treatment with dox for 5 days. Probing for β-ACTIN was used as a protein loading control.

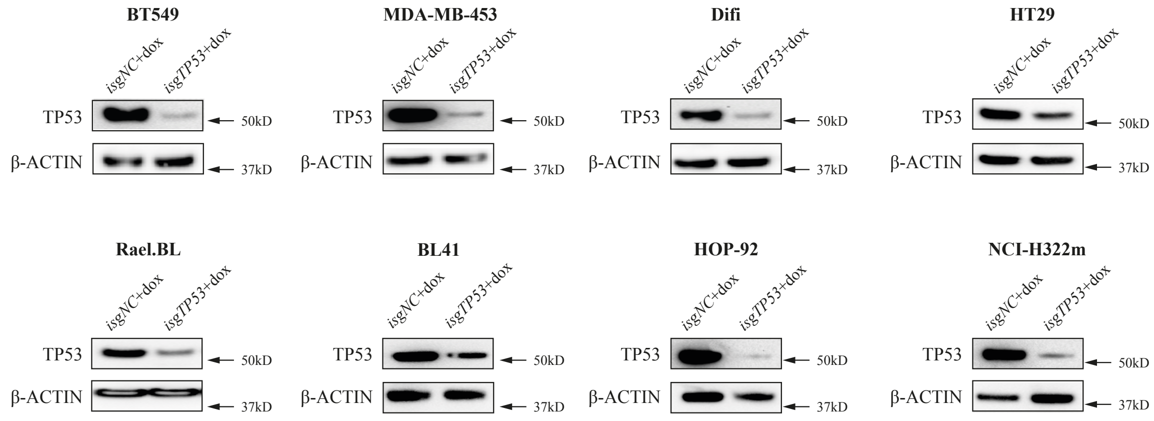

**Extended Data Fig. 2 Loss of mutant TP53 does not impair the sustained survival or impact mitochondrial content and ROS levels in human cancer cell lines**

**a**. *In vitro* survival of the indicated human cancer cell lines with or without dox mediated induction of a mutant *TP53* specific sgRNA or a control sgRNA. **b**. Cell cycle analysis of the cells described in (**a**). **c**. Mitotracker staining of the cells described in (**a**). **d**. CellRox staining of the cells described in (**a**). The analyses described in (**c**) and (**d**) were conducted 2 days after the cells had been treated with dox for 5 days (see (**a**)). Data in (**a**) are presented as mean±SD of experiments conducted in triplicates.

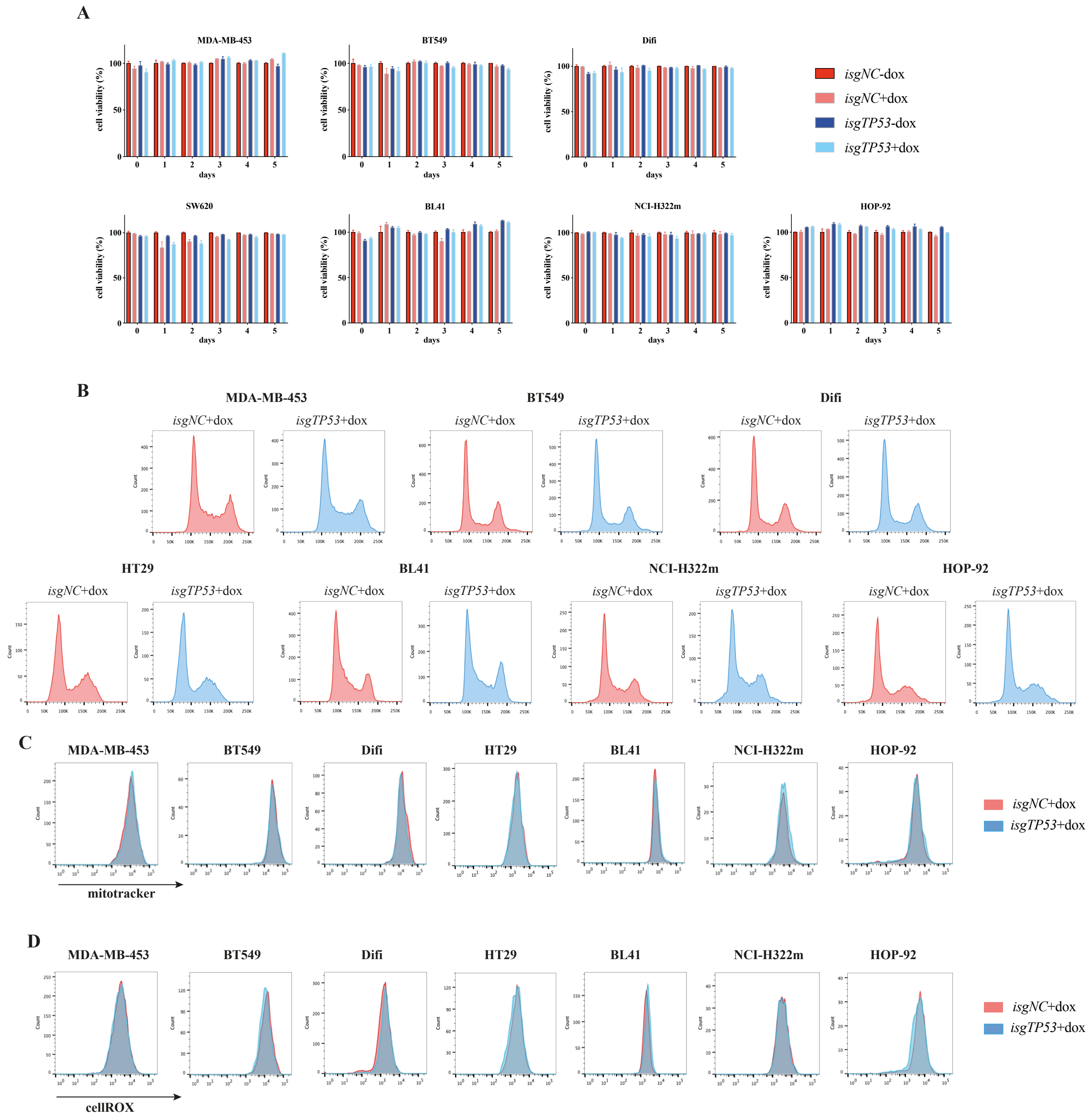

**Extended Data Fig. 3 Loss of mutant TP53 does not impair the ability of human cancer cells to adapt to conditions of stress**

**a**. *In vitro* survival of the indicated human cancer cell lines with or without dox mediated induction of a mutant *TP53* sgRNA or a control sgRNA grown in medium with 1% FCS. **b**. Cell cycle analysis of the cells described in (**a**). **c**. Mitotracker staining of the cells described in (**a**). **d**. CellRox staining of the cells described in (**a**). The analyses described in (**b**), (**c**) and (**d**) were conducted after 2 days in culture in medium with 1% FCS, which was done after the cells had been treated with dox for 5 days in normal medium with 10% FCS. Data in (**a**) are presented as mean±SD of experiments conducted in triplicates.

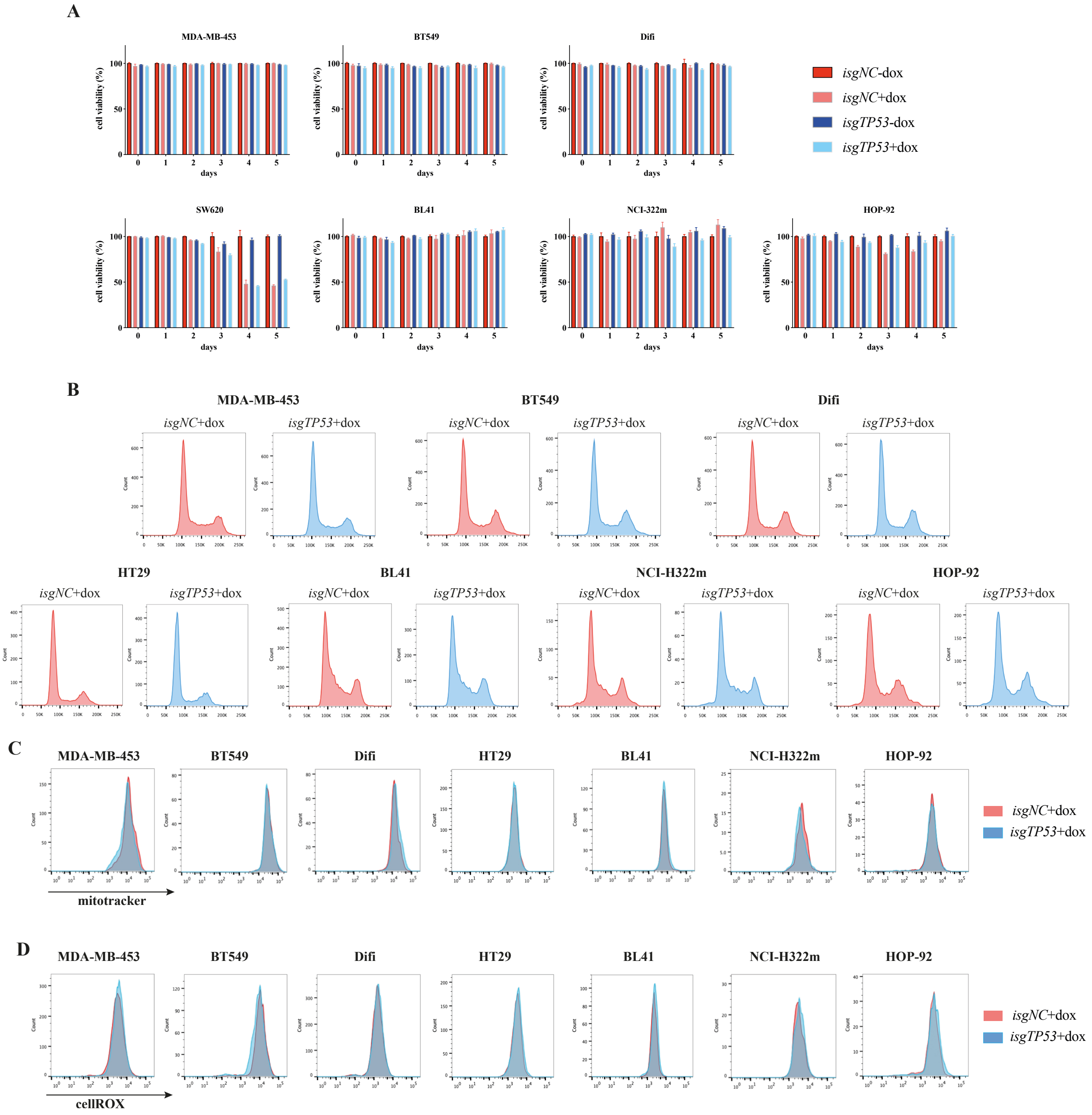

**Extended Data Fig. 4 Loss of mutant TP53 does not impair the sustained proliferation and survival of human cancer cells when cultured in medium containing 3% FCS**

**a**. *In vitro* growth of the indicated human cancer cell lines with or without dox mediated induction of a mutant *TP53* specific sgRNA or a control (NT)sgRNA, grown in medium with 3% FCS. **b**. *In vitro* survival of the cells described in (**a**). Data in (**a**) and (**b**) are presented as mean±SD of experiments conducted in triplicates.

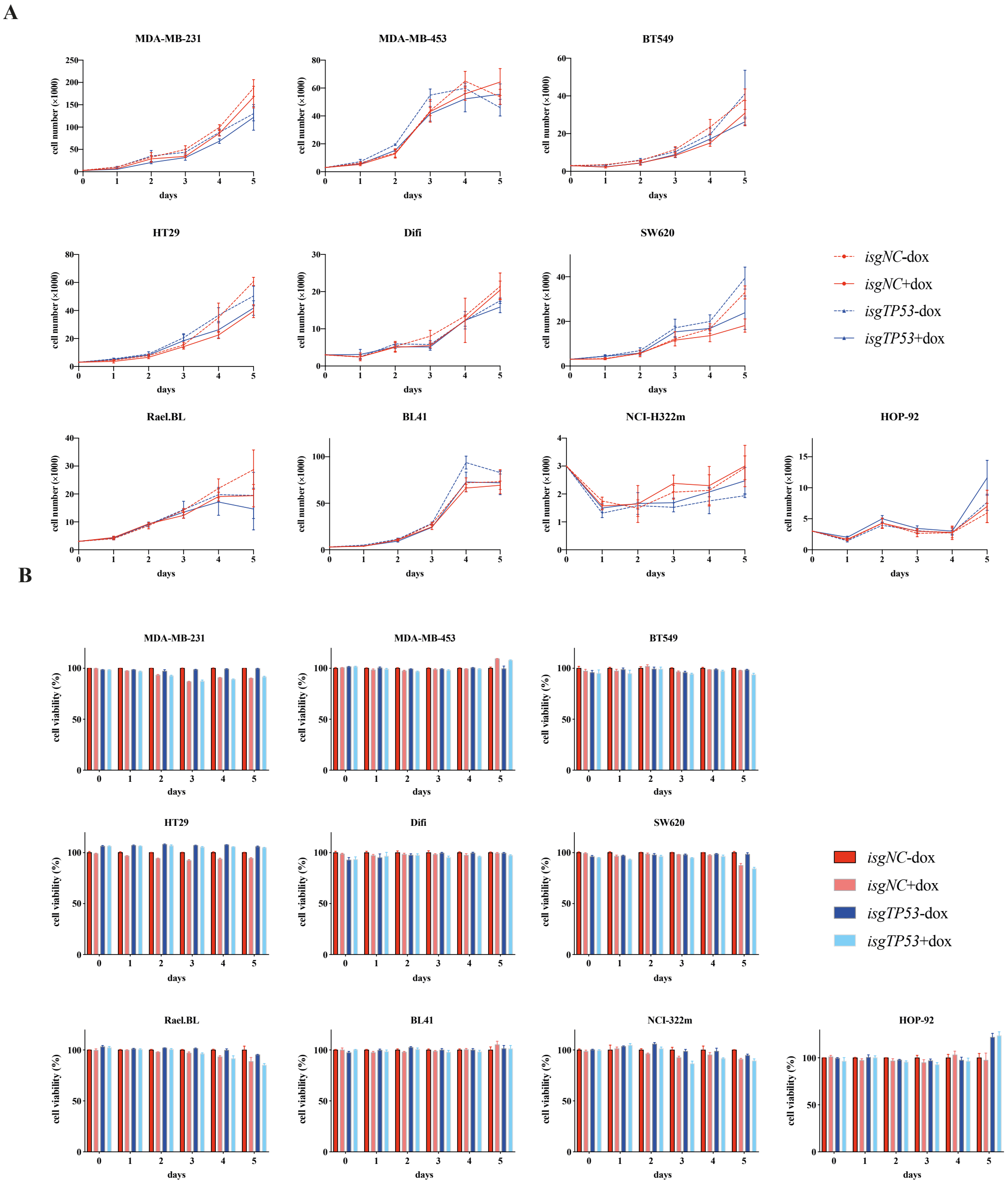

**Extended Data Fig. 5 Loss of mutant TP53 does not impact cell cycling, mitochondrial content or ROS levels in human cancer cell lines cultured in medium with 3% FCS**

**a**. Cell cycle analysis of the indicated human cancer cell lines with or without dox mediated induction of a mutant *TP53* specific sgRNA or a control (NT) sgRNA for 5 days and grown in medium with 3% FCS. **b**. Mitotracker staining of the cells described in (**a**). **c**. CellRox staining of the cells described in (**a**). The analyses described in (**b**) and (**c**) were conducted after 2 days in culture in medium with 3% FCS, which was done after the cells had been treated with dox for 5 days in normal medium with 10% FCS.

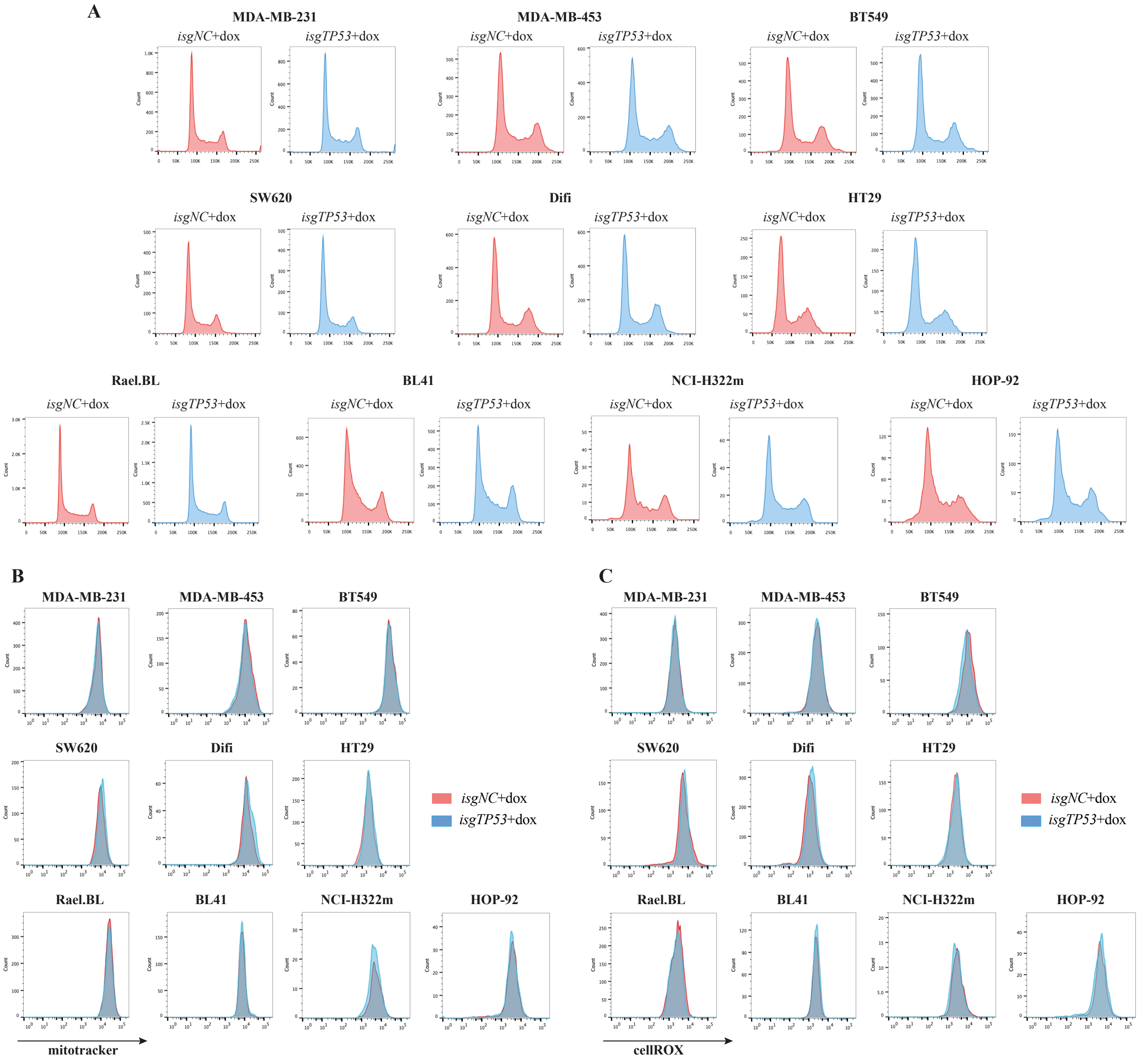

**Extended Data Fig. 6 Loss of mutant p53 does not increase the sensitivity of human cancer cell lines to chemotherapeutic drugs**

The viability of parental mutant TP53 expressing cells and their mutant TP53 depleted derivatives after treatment with the indicated concentrations of etoposide for 48 h was determined by flow cytometric analysis. Data are presented as mean±SD of experiments conducted in triplicates.

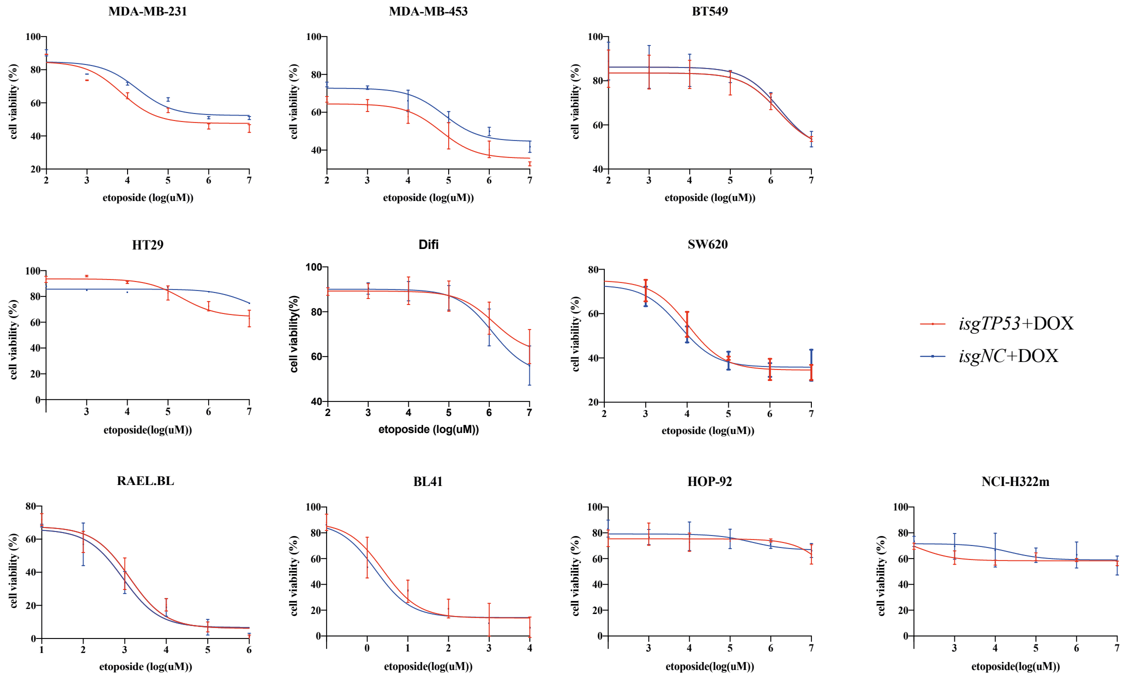

**Extended Data Fig. 7: Single cell clones of MDA-MB-231 cells with complete removal of mutant TP53 do not have impaired survival or proliferation.**

**a**. Western blot analysis showing the complete loss of mutant TP53 in the MDA-MB-231 human breast cancer cell line with a *TP53* specific sgRNA. Probing for β-ACTIN was used as a protein loading control. **b**. *In vitro* growth of parental MDA-MB-231 cells and their mutant TP53 depleted derivatives grown in medium with 1% FCS, 3% FCS or 10% FCS. **c**. *In vitro* survival of the cells described in (**b**). **d**. Cell cycle analysis of the cells described in (**b**). **d**. Mitotracker staining of the cells described in (**b**). **e**. CellRox staining of the cells described in (**b**). **f**. Survival of parental MDA-MB-231 cells and their mutant TP53 depleted derivatives after treatment in culture with the indicated concentrations of etoposide or vehicle for 48 h. Data in (**b**), (**c**) and (**g**) are presented as mean±SD of experiments conducted in triplicates.

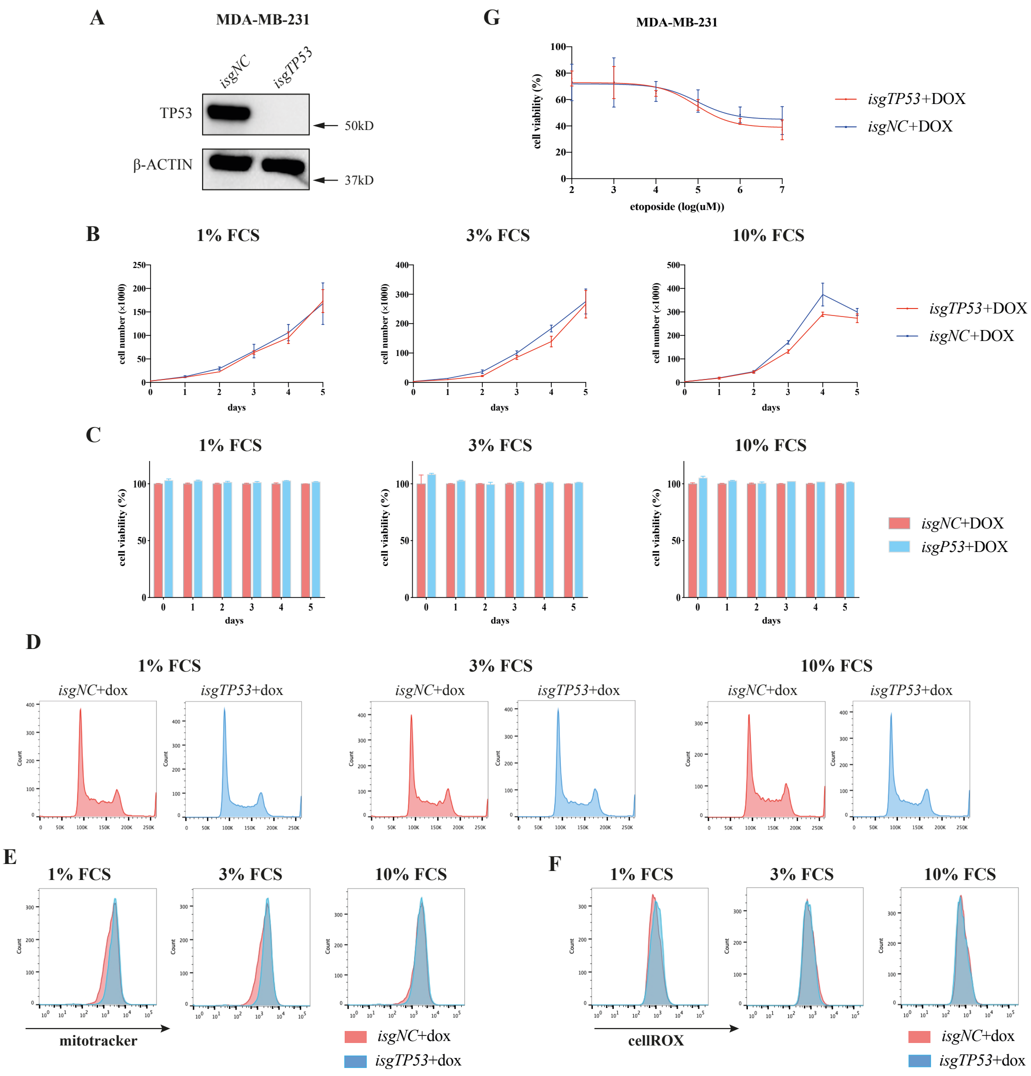

**Extended Data Fig. 8 Complete loss of mutant TP53 in HT29 cells does not impair their sustained survival and proliferation**

**a**. Western blot analysis showing the complete loss of mutant TP53 in single cell cloned HT29 cells with a mutant *TP53* specific sgRNA. Probing for β-ACTIN was used as a protein loading control. **b**. *In vitro* growth of parental cells and their mutant TP53 depleted derivatives grown in medium with 1% FCS, 3% FCS or 10% FCS. **c**. *In vitro* survival of the cells described in (**b**). **d**. Cell cycle analysis of the cells described in (**b**). **d**. Mitotracker staining of the cells described in (**b**). **e**. CellRox staining of the cells described in (**b**). **f**. Survival of parental HT29 cells or their derivatives completely deficient for mutant TP53 after treatment in culture with etoposide or vehicle was determined by flow cytometric analysis. Data in (**b**), (**c**) and (**g**) are presented as mean±SD of experiments conducted in triplicates.

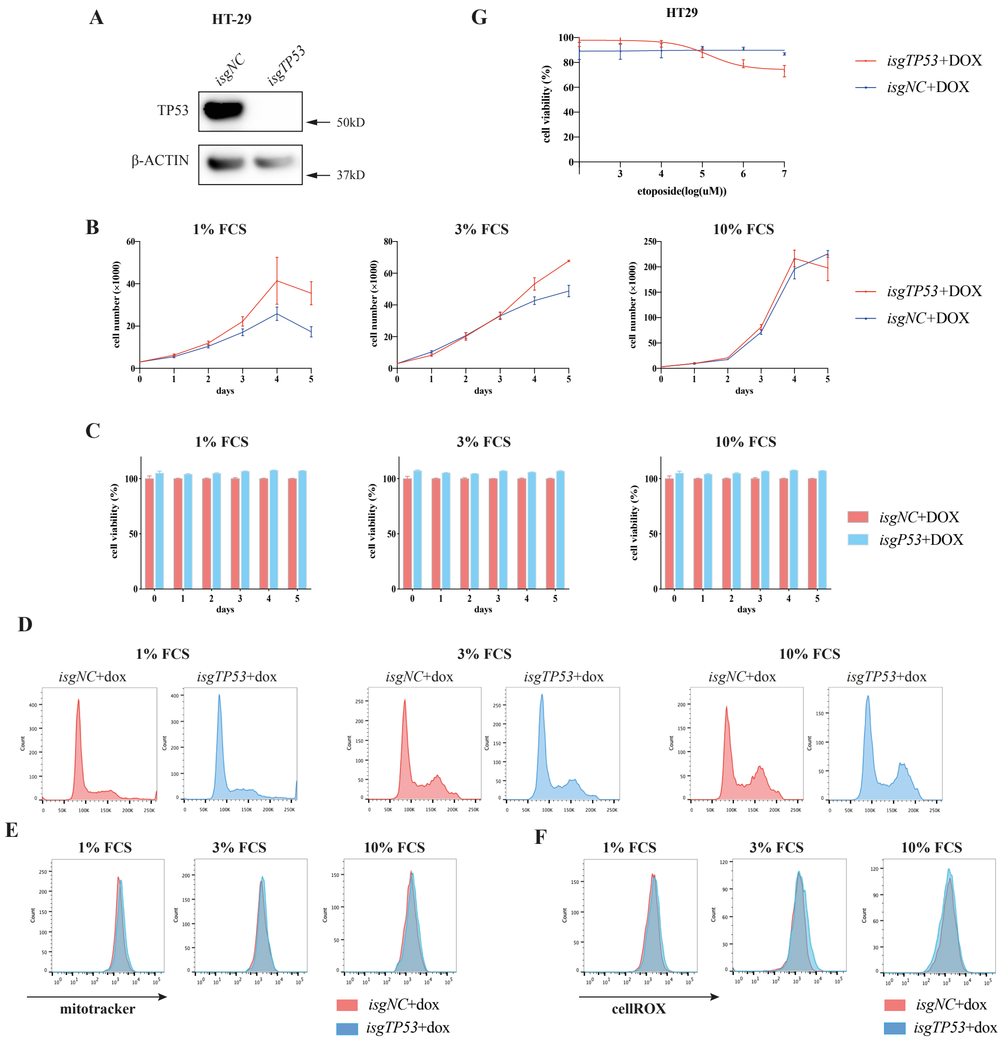

**Extended Data Fig. 9 Complete loss of mutant TP53 in Rael-BL cells does not impair their sustained survival and proliferation**

**a**. Western blot analysis showing the complete loss of mutant TP53 in single cell clones of Rael-BL cells engineered to express a *TP53* specific sgRNA. Probing for β-ACTIN was used as a protein loading control. **b**. *In vitro* growth of parental Rael-BL cells and their mutant TP53-depleted derivatives grown in medium with 1% FCS, 3% FCS or 10% FCS. **c**. *In vitro* survival of the cells described in (**b**) was determined by flow cytometric analysis. **d**. Cell cycle analysis of the cells described in b. **e**. Mitotracker staining of the cells described in (**b**). **f**. CellRox staining of the cells described in (**b**). **g**. Survival of parental Rael-BL cells and their mutant TP53 depleted derivatives after treatment in culture with the indicated concentrations of etoposide or vehicle. Data in (**b**), (**c**) and (**g**) are presented as mean±SD of experiments conducted in triplicates.

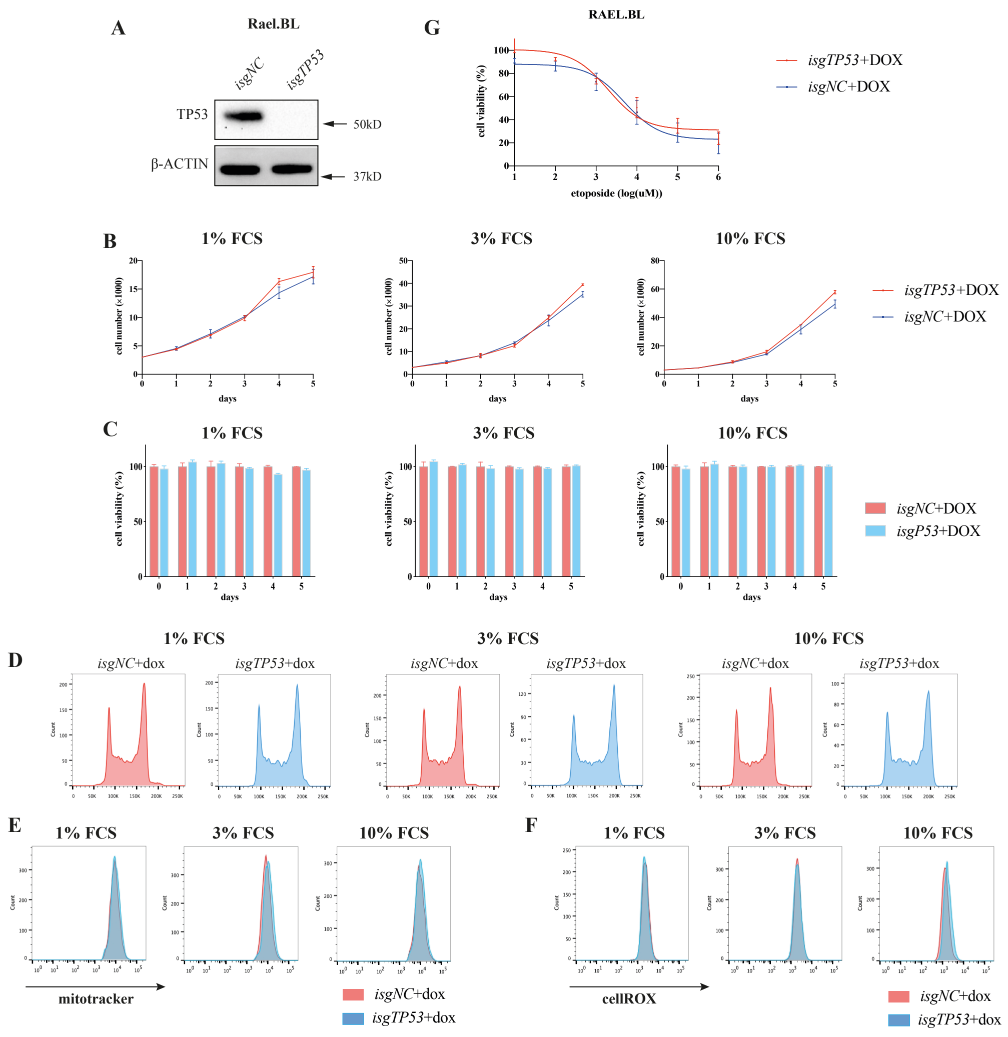

**Extended Data Fig. 10 Flow cytometric analysis of GFP and Ly5.2 expression to established that the malignant cells in recipient mice were derived from the transplanted *Eμ-Myc* lymphoma cells**

*Eμ-Myc* lymphoma cells (C57BL/6J-Ly5.2 background) were identified in the spleens of recipient (tumour cell transplanted) mice (Ly5.1 background) and as GFP and Ly5.2 double positive.

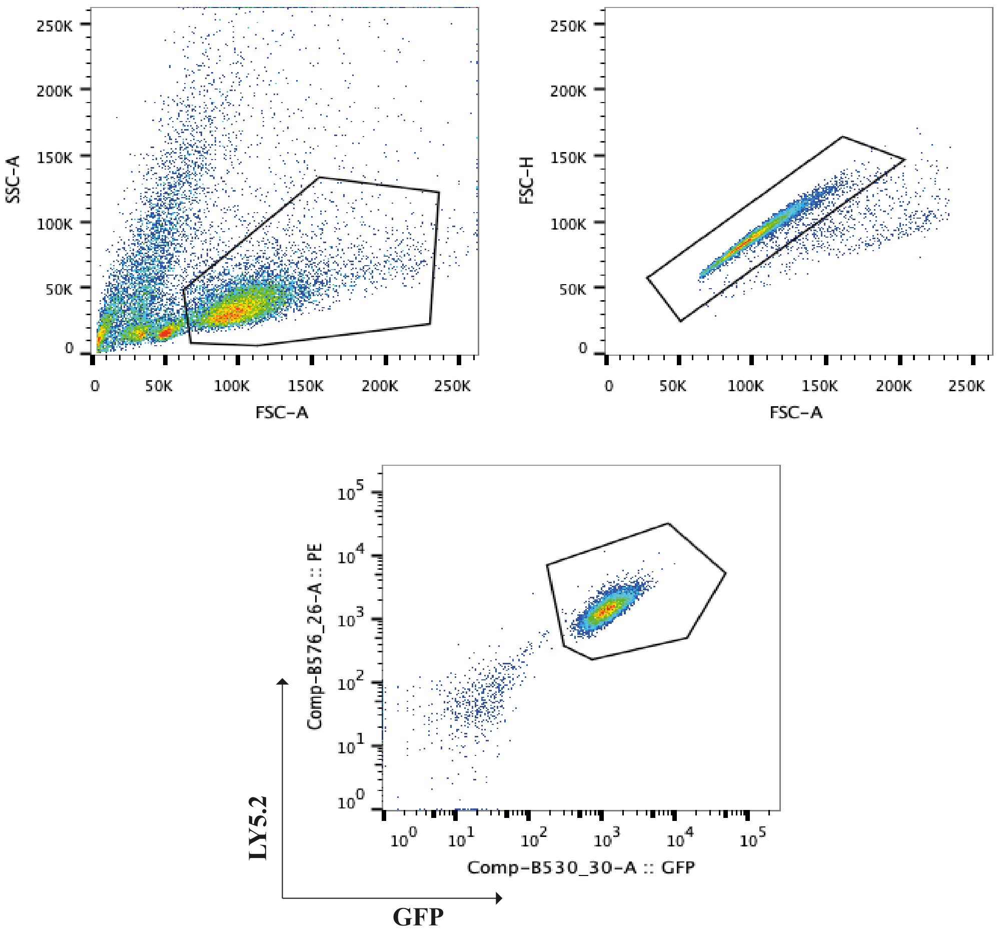

**Extended Data Fig. 11 Impact of the loss of mutant TP53 caused by RNA interference on cell survival and cell proliferation**

**a**. Western blot analysis showing the reduction of mutant TP53 protein in the indicated human cancer cell lines with or without dox mediated induction of a *TP53* specific shRNA or a control shRNA. Probing for β-ACTIN was used as a protein loading control. **b**. *In vitro* growth of the cells with or without dox mediated induction of a *TP53* specific shRNA or a control shRNA for 5 days. **c**. *In vitro* survival of the cells described in (**b**). Data in (**b**) and (**c**) are presented as mean±SD of experiments conducted in triplicates.

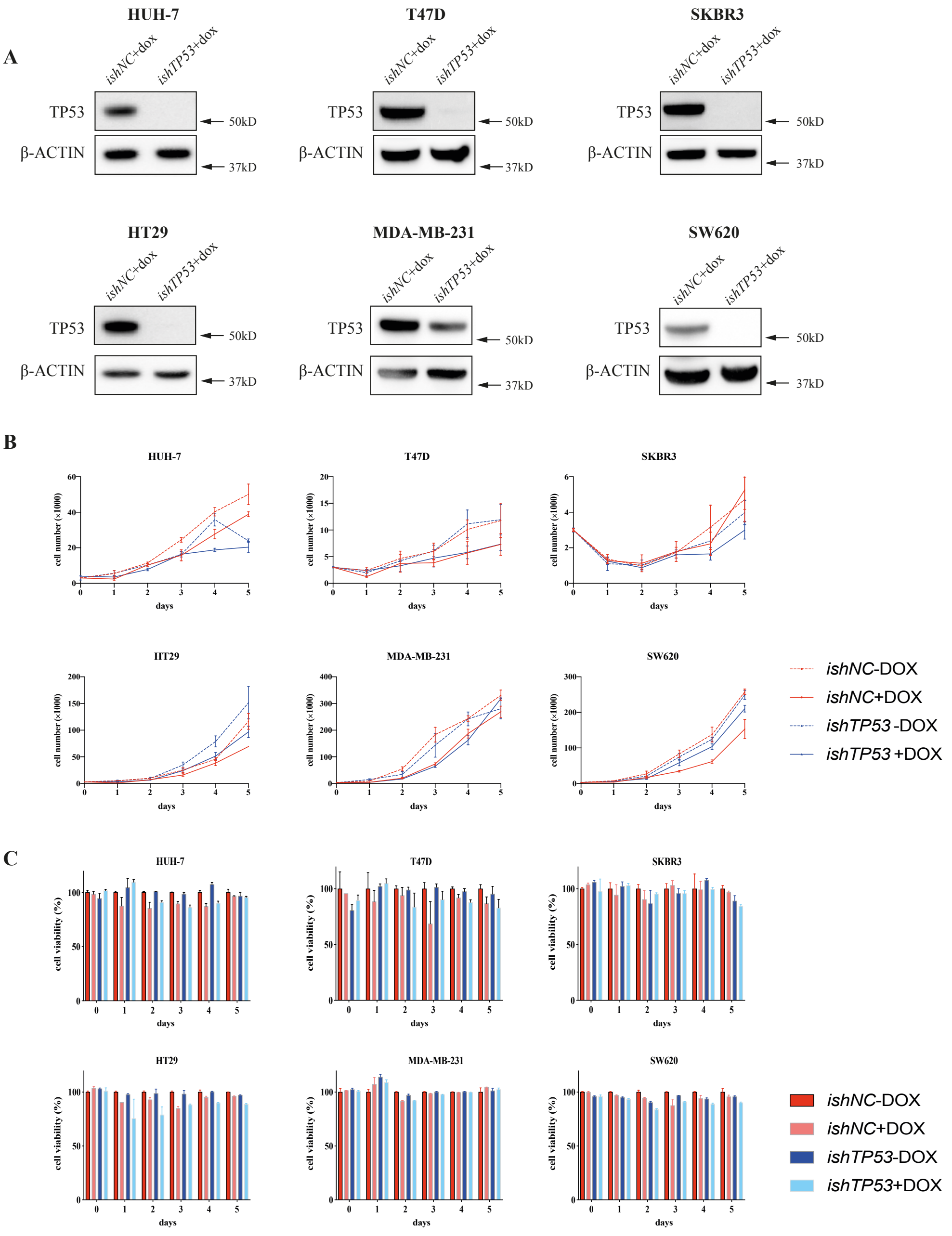

**Extended Data Table 1 Statistical analysis of differences in the numbers of cells in culture in medium containing 1%, 3% or 10% FCS comparing cells with the mutant *TP53* specific sgRNA *vs* cells with a control sgRNA after dox treatment**

Cell numbers in culture were determined from day 1 to day 5 of dox treatment. Data are presented as mean±SD of experiments conducted in triplicates. Statistical significance of differences between the cells with the mutant TP53 specific sgRNA vs the cells with the control sgRNA was determined by using GraphPad.

### MDA-MB-231

| | 1% FCS | | | 3% FCS | | | 10% FCS | | |
|---|---|---|---|---|---|---|---|---|---|
| | ***isgNC* +dox** | ***isgTP53* +dox** | ***p* value** | ***isgNC*+dox** | ***isgTP53* +dox** | ***p* value** | ***isgNC*+dox** | ***isgTP53* +dox** | ***p* value** |
| **D1** | 4.206±0.481 | 4.084±0.224 | 0.710980183 | 7.235±0.682 | 5.533±0.303 | 0.016858348 | 6.305±0.075 | 6.117±1.298 | 0.814240594 |
| **D2** | 11.446±0.571 | 10.425±0.141 | 0.03981823 | 28.886±7.559 | 20.668±2.730 | 0.151281458 | 22.811±1.062 | 21.520±1.943 | 0.369699861 |
| **D3** | 24.328±1.914 | 21.834±1.243 | 0.131349171 | 34.392±2.833 | 31.677±6.042 | 0.519908632 | 42.166±5.401 | 46.143±3.087 | 0.330357148 |
| **D4** | 50.134±2.655 | 49.768±10.276 | 0.955246685 | 86.221±6.132 | 68.362±5.089 | 0.017818634 | 130.265±17.452 | 126.77±12.416 | 0.354568118 |
| **D5** | 75.873±1.403 | 69.148±11.251 | 0.362385913 | 167.129±24.138 | 121.726±28.898 | 0.105009889 | 247.305±55.493 | 224.454±19.346 | 0.537569021 |

### MDA-MB-453

| | 1% FCS | | | 3% FCS | | | 10% FCS | | |
|---|---|---|---|---|---|---|---|---|---|
| | ***isgNC*+dox** | ***isgTP53* +dox** | ***p* value** | ***isgNC*+dox** | ***isgTP53* +dox** | ***p* value** | ***isgNC*+dox** | ***isgTP53* +dox** | ***p* value** |
| **D1** | 2.386±0.084 | 3.151±0.717 | 0.140628776 | 5.262±0.871 | 5.542±0.434 | 0.645225083 | 4.799±0.259 | 4.892±0.219 | 0.658763828 |
| **D2** | 5.918±0.84 | 7.414±1.072 | 0.129902051 | 12.82±3.003 | 15.315±0.636 | 0.231916555 | 15.825±1.383 | 13.028±0.480 | 0.029693958 |
| **D3** | 13.556±1.505 | 14.358±1.882 | 0.595458952 | 43.097±2.939 | 41.549±5.293 | 0.680904864 | 23.844±3.365 | 18.844±0.695 | 0.065343274 |
| **D4** | 22.88±4.660 | 22.769±1.104 | 0.969760573 | 55.868±4.765 | 52.242±9.270 | 0.579285999 | 54.220±4.1 | 57.550±1.988 | 0.274289253 |
| **D5** | 37.897±0.736 | 40.66±11.123 | 0.689881175 | 64.28±9.613 | 55.552±7.417 | 0.281093082 | 44.652±7.098 | 48.662±4.128 | 0.445204854 |

## BT549

| | 1% FCS | | | 3% FCS | | | 10% FCS | | |
|---|---|---|---|---|---|---|---|---|---|
| | ***isgNC* +dox** | ***isgTP53* +dox** | ***p* value** | ***isgNC*+dox** | ***isgTP53* +dox** | ***p* value** | ***isgNC*+dox** | ***isgTP53* +dox** | ***p* value** |
| **D1** | 2.023±0.367 | 1.737±0.622 | 0.53017328 | 2.288±0.517 | 2.379±0.230 | 0.792799421 | 2.971±0.667 | 3.107±0.012 | 0.741597686 |
| **D2** | 3.15±0.224 | 2.741±0.259 | 0.227522664 | 4.47±1.181 | 4.358±0.685 | 0.893200313 | 5.45±1.231 | 6.488±1.212 | 0.654475522 |
| **D3** | 5.676±0.051 | 5.448±0.268 | 0.222470301 | 8.425±0.975 | 8.945±1.669 | 0.665598324 | 9.684±1.063 | 9.292±0.677 | 0.618479407 |
| **D4** | 7.808±0.492 | 6.342±1.415 | 0.165423405 | 15.053±1.708 | 17.159±2.627 | 0.309194169 | 23.852±2.525 | 31.704±1.251 | 0.008488246 |
| **D5** | 12.919±2.79 | 11.42±1.577 | 0.463281261 | 30.86±6.697 | 26.251±1.677 | 0.31192849 | 47.71±10.285 | 54.28±5.341 | 0.381801245 |

## HT29

| | 1% FCS | | | 3% FCS | | | 10% FCS | | |
|---|---|---|---|---|---|---|---|---|---|
| | ***isgNC*+dox** | ***isgTP53*+dox** | ***p* value** | ***isgNC*+dox** | ***isgTP53*+dox** | ***p* value** | ***isgNC*+dox** | ***isgTP53*+dox** | ***p* value** |
| **D1** | 2.326±0.13 | 2.893±0.298 | 0.039434729 | 3.629±1.038 | 4.943±0.970 | 0.1846181 | 2.282±2.19 | 2.017±0.112 | 0.135892499 |
| **D2** | 3.744±0.27 | 4.04±0.735 | 0.548457658 | 6.558±1.012 | 8.239±0.399 | 0.055536048 | 5.238±0.676 | 6.688±1.272 | 0.753783072 |
| **D3** | 8.835±0.523 | 8.763±0.824 | 0.90438818 | 14.193±1.748 | 18.463±4.932 | 0.230448431 | 22.633±6.123 | 30.820±6.180 | 0.178442211 |
| **D4** | 13.852±0.836 | 14.117±1.981 | 0.841598161 | 22.519±2.571 | 26.201±5.987 | 0.383231937 | 67.926±10.853 | 76.093±13.901 | 0.467449016 |
| **D5** | 12.94±1.319 | 13.799±1.973 | 0.564485477 | 39.469±4.629 | 41.796±5.252 | 0.595686942 | 64.293±10.916 | 63.648±17.671 | 0.959704736 |

## SW620

| | 1% FCS | | | 3% FCS | | | 10% FCS | | |
|---|---|---|---|---|---|---|---|---|---|
| | ***isgNC*+dox** | ***isgTP53*+dox** | ***p* value** | ***isgNC*+dox** | ***isgTP53*+dox** | ***p* value** | ***isgNC*+dox** | ***isgTP53*+dox** | ***p* value** |
| **D1** | 3.077±0.407 | 3.431±0.346 | 0.315513166 | 3.207±0.385 | 4.404±0.679 | 0.056712845 | 2.575±0.523 | 3.392±0.256 | 0.072291708 |
| **D2** | 4.883±0.635 | 3.589±0.215 | 0.028818835 | 5.686±1.492 | 5.771±0.953 | 0.937896372 | 5.355±2.214 | 6.106±1.194 | 0.621804321 |
| **D3** | 2.131±0.52 | 1.847±0.328 | 0.469311904 | 11.621±0.464 | 15.33±2.559 | 0.068949389 | 16.884±4.493 | 17.855±2.45 | 0.758990716 |
| **D4** | 1.017±0.284 | 0.867±0.09 | 0.435418852 | 13.6±2.692 | 16.843±1.044 | 0.123640931 | 53.177±11.008 | 68.367±22.223 | 0.348576212 |
| **D5** | 1.144±0.173 | 1.198±0.087 | 0.654418352 | 18.148±2.965 | 23.884±7.526 | 0.286700363 | 119.219±12.036 | 192.295±11.161 | 0.001471749 |

### Difi

| | 1% FCS | | | 3% FCS | | | 10% FCS | | |
|---|---|---|---|---|---|---|---|---|---|
| | ***isgNC*+dox** | ***isgTP53*+dox** | ***p* value** | ***isgNC*+dox** | ***isgTP53*+dox** | ***p* value** | ***isgNC*+dox** | ***isgTP53*+dox** | ***p* value** |
| **D1** | 3.062±0.793 | 2.968±0.187 | 0.851809315 | 2.457±1.062 | 3.124±1.368 | 0.541371681 | 2.962±0.300 | 3.092±0.612 | 0.75948847 |
| **D2** | 4.241±0.537 | 4.114±0.637 | 0.805384018 | 5.027±1.302 | 5.198±0.487 | 0.841983214 | 7.199±0.876 | 6.132±0.977 | 0.232408569 |
| **D3** | 10.442±1.331 | 11.296±0.660 | 0.376101124 | 5.484±0.567 | 5.134±0.874 | 0.591856893 | 15.838±0.699 | 10.207±1.175 | 0.002044333 |
| **D4** | 12.649±0.906 | 11.927±1.441 | 0.50334788 | 12.296±5.958 | 12.301±2.358 | 0.99910596 | 28.186±1.058 | 28.285±3.888 | 0.968060156 |
| **D5** | 19.493±4.305 | 16.112±2.160 | 0.291068384 | 20.416±2.425 | 15.852±1.545 | 0.051462534 | 47.065±1.96 | 48.907±6.017 | 0.640736306 |

### Rael.BL

| | 1% FCS | | | 3% FCS | | | 10% FCS | | |
|---|---|---|---|---|---|---|---|---|---|
| | ***isgNC*+dox** | ***isgTP53*+dox** | ***p* value** | ***isgNC*+dox** | ***isgTP53*+dox** | ***p* value** | ***isgNC*+dox** | ***isgTP53*+dox** | ***p* value** |
| **D1** | 3.560±0.318 | 3.450±0.218 | 0.646201948 | 4.404±0.472 | 4.177±0.429 | 0.570702638 | 4.243±0.358 | 4.997±0.514 | 0.106109061 |
| **D2** | 4.711±0.647 | 5.258±0.296 | 0.253947839 | 9.319±0.102 | 9.147±0.193 | 0.247098259 | 7.082±0.513 | 7.761±0.58 | 0.203236938 |
| **D3** | 7.631±0.953 | 8.908±0.543 | 0.113929531 | 12.3±1.047 | 13.498±0.998 | 0.224910888 | 14.999±2.452 | 12.519±2.63 | 0.298266439 |
| **D4** | 10.542±0.701 | 15.267±1.506 | 0.007899692 | 19.088±1.569 | 17.111±4.783 | 0.533822361 | 24.934±4.021 | 19.638±2.086 | 0.112933236 |
| **D5** | 11.213±0.947 | 14.487±1.453 | 0.071096083 | 19.374±3.996 | 14.629±7.479 | 0.387354741 | 21.592±7.193 | 19.39±3.523 | 0.658827305 |

## BL41

| | 1% FCS | | | 3% FCS | | | 10% FCS | | |
|---|---|---|---|---|---|---|---|---|---|
| | ***isgNC*+dox** | ***isgTP53*+dox** | ***p* value** | ***isgNC*+dox** | ***isgTP53*+dox** | ***p* value** | ***isgNC*+dox** | ***isgTP53*+dox** | ***p* value** |
| **D1** | 3.503±0.311 | 3.47±0.099 | 0.869569274 | 3.756±0.138 | 4.097±0.142 | 0.04108204 | 4.718±0.359 | 3.947±0.586 | 0.124280328 |
| **D2** | 5.662±0.251 | 6.89±1.058 | 0.122147139 | 10.635±1.284 | 9.246±0.875 | 0.196727687 | 6.507±0.239 | 5.334±0.173 | 0.002360913 |
| **D3** | 12.647±1.598 | 12.883±0.466 | 0.935578191 | 24.656±2.913 | 24.194±0.482 | 0.799484497 | 15.726±1.113 | 15.225±1.469 | 0.664293937 |
| **D4** | 17.311±3.243 | 19.254±2.1 | 0.43290805 | 66.302±4.05 | 72.857±10.565 | 0.372485437 | 44.080±5.176 | 40.858±5.497 | 0.501013093 |
| **D5** | 19.991±6.21 | 20.588±4.445 | 0.898716193 | 69.333±4.543 | 72.075±12.822 | 0.744619422 | 150.922±9.915 | 112.393±40.369 | 0.183686247 |

### NCI-H322m

| | 1% FCS | | | 3% FCS | | | 10% FCS | | |
|---|---|---|---|---|---|---|---|---|---|
| | ***isgNC*+dox** | ***isgTP53*+dox** | ***p* value** | ***isgNC*+dox** | ***isgTP53*+dox** | ***p* value** | ***isgNC*+dox** | ***isgTP53*+dox** | ***p* value** |
| **D1** | 2.771±0.287 | 2.424±0.307 | 0.227517358 | 1.571±0.141 | 1.5±0.154 | 0.592980796 | 1.454±0.05 | 1.534±0.213 | 0.563416996 |
| **D2** | 2.529±0.959 | 2.681±0.663 | 0.833026737 | 1.643±0.663 | 1.658±0.392 | 0.974114871 | 2.363±0.475 | 2.054±0.410 | 0.443165967 |
| **D3** | 2.578±0.37 | 2.392±0.693 | 0.703066035 | 2.379±0.296 | 1.689±0.164 | 0.024380876 | 2.999±0.449 | 2.906±0.556 | 0.833429768 |
| **D4** | 3.94±0.808 | 3.535±0.598 | 0.524703368 | 2.3±0.688 | 2.072±0.174 | 0.607398696 | 4.253±0.282 | 3.731±0.659 | 0.275757177 |
| **D5** | 3.014±0.406 | 2.76±0.248 | 0.406962635 | 3.003±0.741 | 2.472±0.482 | 0.356702252 | 4.71±0.762 | 4.443±0.763 | 0.691067585 |

**HOP-92**

| | 1% FCS | | | 3% FCS | | | 10% FCS | | |
|---|---|---|---|---|---|---|---|---|---|
| | ***isgNC*+dox** | ***isgTP53*+dox** | ***p* value** | ***isgNC*+dox** | ***isgTP53*+dox** | ***p* value** | ***isgNC*+dox** | ***isgTP53*+dox** | ***p* value** |
| **D1** | 1.829±0.326 | 1.608±0.393 | 0.495929201 | 1.691±0.222 | 2.019±0.171 | 0.113412843 | 1.519±0.042 | 2.084±0.116 | 0.023265387 |
| **D2** | 1.49±0.27 | 1.501±0.243 | 0.964033627 | 4.299±0.788 | 5.007±0.537 | 0.268078965 | 1.884±0.366 | 2.274±0.21 | 0.278005574 |
| **D3** | 2.231±0.314 | 5.796±0.085 | 0.19037752 | 2.985±0.425 | 3.439±0.418 | 0.258056518 | 4.371±0.338 | 5.137±0.757 | 0.219294532 |
| **D4** | 2.499±0.426 | 2.712±0.273 | 0.506640321 | 2.8±0.863 | 3.004±0.481 | 0.739176303 | 6.808±0.388 | 9.04±0.113 | 0.000674358 |
| **D5** | 2.084±0.198 | 2.331±0.114 | 0.136082767 | 6.997±2.613 | 11.595±2.829 | 0.107522706 | 7.288±0.752 | 9.518±1.384 | 0.070379688 |
